# Emerging Haemagglutinin Mutations in Bovine-origin H5N1 Influenza Viruses from Humans and Cattle Retain Avian Receptor Binding with Increased Stability

**DOI:** 10.64898/2026.08.18.745528

**Authors:** Jiayun Yang, Thomas P. Peacock, Kristel Ramirez Valdez, Jie Zhou, Hannah J. Klim, Ksenia Sukhova, Jean-Remy Sadeyen, Ian H. Brown, Wendy S. Barclay, Munir Iqbal

## Abstract

The current H5N1 panzootic has seen an unprecedented host range expansion, including sustained circulation in US dairy cattle, detected in March 2024. By July 2026, infections had been reported on more than 1,150 dairy farms across 19 states. Although the outbreak initially centred in Texas, California has emerged as the principal focus of transmission and accounts for most human infections associated with exposure to infected dairy cattle. Continued transmission in cattle and repeated spillover into humans increase opportunities for acquisition of mammalian-adaptive mutations that could elevate zoonotic and pandemic risk.

The haemagglutinin (HA) protein plays a central role in modulating virus receptor binding and airborne transmission. Here, we characterised the receptor-binding and stability phenotypes of HA mutations identified in viruses circulating in Californian dairy cattle. Receptor-binding specificity was assessed using bio-layer interferometry and pseudotype virus entry assays. All tested HA variants maintained a preference for avian-type α2,3-linked sialic acid receptors. We evaluated HA stability using fusion and thermostability assays. All mutants exhibited fusion pH values >5.5, outside the range associated with efficient airborne transmission in humans (pH 5.0-5.5). However, mutations D88G and S94N increased pH stability, with fusion pH values of 5.6 and 5.7, respectively, compared with 5.9 for wild-type HA. Viruses harbouring both mutations displayed increased thermostability.

These findings demonstrate that cattle-origin H5N1 viruses retain avian-like receptor specificity despite acquiring mutations that modestly enhance HA stability. Evolution of H5N1 viruses in dairy cattle underscores the importance of genomic and phenotypic surveillance to identify mutations that may increase zoonotic risk.

## Introduction

Since the first detection of H5N1 high pathogenicity avian influenza virus (HPAIV) in 1996 in Guangdong, China, the A/goose/Guangdong/1/96 (Gs/Gd/96) HPAIV has rapidly evolved and diversified into multiple clades. Among the many clades that have emerged, clade 2.3.4.4 has been particularly persistent in avian species since 2010, the clade then subsequently diversified into clade 2.3.4.4a through to 2.3.4.4h (1). The clade 2.3.4.4b H5N1 virus has become the most widespread and persistent since 2020, resulting in the current panzootic causing significant mortality of wild animals and economic losses to the agricultural sector (2).

The current clade 2.3.4.4b H5N1 viruses have a much-expanded host range compared to previous H5 HPAIVs of other clades, with detections in multiple orders of wild birds, domestic poultry, carnivorous mammals including mink, foxes, domestic cats, and marine mammals (3–6). The virus further expanded its host range with detections in dairy cows, firstly reported in Kansas and Texas on 25^th^ March 2024, in the USA (7). Since then (to March 2026), the virus has rapidly spread to 19 states, affecting over 1,106 cattle farms according to the reports the United States Department of Agriculture (USDA) (8). The high density of dairy cattle in California subsequently resulted in this becoming the second epicentre of the outbreak (9). The first detection of H5N1 in cattle in California was reported in August 2024. Since then, the virus has been widely detected in California, with the total number of detections dwarfing those in the other 20 states with 773 reported farms in California. The US Centres for Disease Control and Prevention (CDC) has confirmed 71 human infections with the clade 2.3.4.4b H5N1 virus. Forty-one of the human cases were linked to exposure to dairy cattle, and 36 of the 41 cases are from California (10). The increasing numbers of infections in dairy cattle and humans suggests heightened zoonotic opportunity for the H5N1 virus to spillover to humans. To date however these viruses have lacked the capacity to spread between humans (11).

The receptor binding of influenza A virus (IAV) to host cell sialic acid (SA) is critical in virus infection. Avian influenza viruses tend to have binding preference to glycans containing α2,3-(SAα-2,3Gal)-linked SA, whereas human influenza viruses preferentially bind to α2,6- (SAα-2,6Gal)-linked SA (12). The consequence of viral glycoprotein haemagglutinin (HA) binding to SA of different linkages can change or expand virus host range and tissue tropism. Both human α2,6-linked SA and avian α2,3-linked SA have been detected in bovine mammary glands (13, 14), suggesting cattle as a novel intermediate host for avian influenza virus (AIV), to acquire adaptive mutations to cross the host barrier from avian to mammalian species.

In addition, to receptor binding, viral HA undergoes pH-dependent conformational changes to mediate membrane fusion during entry into host cells. The HA of human transmissible influenza viruses fuse at a lower pH ≤5.5, whereas HA of avian influenza viruses are usually less stable, fusing above pH 5.5 (15, 16). The pH stability is linked to the airborne transmission routes of human influenza viruses requiring more stable HA to survive environmental conditions found in aerosol particles and acidic respiratory fluids. HA thermostability has also been identified as a key factor in transmission of AIV and has often been shown to correlate strongly with pH stability (17).

In this study, we investigated the HA properties of the emerging bovine H5N1 mutants associated with the epicentre of California, in their receptor binding affinity, virus entry preference, thermostability and pH stability. These HA functions are key molecular determinants of host adaptation and transmissibility. Our data provide timely and critical insights to monitor the zoonotic risk of these bovine H5N1 viruses, informing ongoing surveillance efforts and assessments quantifying relative risks for human infections.

## Materials and Methods

### Ethics Statement

The experimental procedures involving embryonated chicken eggs were undertaken in strict accordance with the guidance and regulations of the UK Home Office under the licence number PP671846. The work was scrutinised and approved by the animal welfare ethical review board at The Pirbright Institute. Incorporating the 3Rs and incorporating ARRIVE (Animal Research: Reporting of *in vivo* experiment) guidelines for quality, reproducibility, and translatability of animal studies.

### Cells and Viruses

Human embryonic kidney 293T (HEK-293T) cells, the parental Madin-Darby canine kidney (MDCK) cells and African green monkey kidney epithelial (Vero) cells were acquired from the Central Service Unit (CSU) of The Pirbright Institute (TPI). HEK293 cells lacking sialyltransferase activities (HEK293 ΔST3GAL1/2/3/4/5/6 ΔST61/2; aka 293delST cells) were a kind gift from Professors Henrik Clausen and Yoshiki Narimatsu from the University of Copenhagen (18, 19). The cell lines were cultured by Dulbecco’s Modified Eagle’s medium (DMEM) and supplemented with 10% fetal bovine serum (FBS) (Gibco) with 5% CO_2_ at 37°C. The HA and NA sequences of A/cattle/Texas/24-008749-003/2024 (H5N1) (NCBI accession PP755589.1 and PP755591.1, respectively), referred to as TX-Cattle WT, and A/cattle/CA/24-036360-002-original/2024(H5N1) (NCBI accession PQ829557.1 and PQ829559.1, respectively), referred to as CA-Cattle WT, were acquired from NCBI and synthesised by GeneScript and cloned into pHW2000 vector. The cleavage site of H5 HA has been replaced by a monobasic cleavage site as previously described (20). Virus rescues were performed as previously described using internal segments from the laboratory adapted strain A/Puerto Rico/8/1934 (H1N1) (PR8). The rescued viruses including TX-Cattle WT, CA- Cattle WT, and HA mutant viruses based on TX-Cattle WT HA, D88G, S94N, V131A, V131E, and V131M, were propagated in 9- to 10-day old embryonated chicken eggs.

### Site-directed mutagenesis (SDM)

HA plasmids with targeted mutations were generated using single-site mutagenesis QuikChange II kit (Agilent) following manufacturer’s instructions.

### Bio-layer interferometry (BLI)

BLI was performed as previously described. In brief, each rescued virus was propagated in 9- to 10-day old embryonated chicken embryos, the propagated virus was then purified through 30% and 60% sucrose gradient at 27,000 rpm for 2h at 4 . The purified viruses were normalised to 100pmol by quantifying viral nucleoprotein (NP) expression using enzyme-linked immunosorbent assay (ELISA) as previously described (21). The standardised viruses were then diluted with 10 μM oseltamivir carboxylate (Roche) and 10 μM zanamivir (GSK) against avian-like sugar analogue 3SLN and human-like sugar analogue 6SLN, and receptor binding affinity was tested by Octet® R8 system (Sartorius) using streptavidin biosensors (Sartorius). The virus binding to each sugar analogue was normalised to fractional saturation and the concentrations of sugar loading as previously described (22).

### Syncytium formation assays

Monolayered Vero cells were infected with 1.0 multiplicity of infection (MOI) of each virus for 1h at 37°C. The infected cells were then washed with PBS for three times and incubated at 37°C for 15h. The cells were further treated with 3.0 μg/mL N-tosyl-L-phenylalanine chloromethyl ketone (TPCK) trypsin in DMEM for 15min and followed by incubation with PBS ranging from pH 5.2 to 6.0 at 0.1 pH increment for 5min. The Vero cells were then maintained with DMEM with 10% FBS for 3h. The cells were then fixed with acetone: methanol (1:1 ratio) and stained with 20% Giemsa solution (Sigma-Aldrich). Stained cell images were taken by EVOS XL imaging system at 400µm (Life Technologies). The lowest pH that induced visible syncytium formation is recorded as the pH of fusion with multinucleated cells and indistinct cell membranes.

### Thermostability assay

Briefly, the tested viruses were standardised to 32HAU/50 µL, and then incubated in thermal cycler (Bio-Rad) at 50°C, 50.7°C, 51.9°C, 53.8°C, 56.1°C, 58.0°C, 59.2°C, and 60°C and 4°C as control for 30min, the HA titres were then determined by Haemagglutination assay using 1% chicken red blood cells (cRBCs).

### Generation of pseudoviruses

The VSV-G expression plasmid, and lentiviral packaging genes and luciferase genome were used as previously described (23). The following human codon optimised HA, NA, protease and sialyl-transferase expression plasmids were synthesised by Genscript in pcDNA3.1: H1HA - A/Rhode Island/04/2016 (H1N1pdm09; ANM90381.1), N1NA - A/England/195/2009 (H1N1pdm09; ACR15618.1), H7HA and N9NA - A/Shanghai/02/2013 (H7N9; YP_009118475.1, YP_009118481.1), H5HA - A/cattle/Texas/24-008749-003/2024 (H5N1; PP755589.1), H5HA – A/cattle/CA/24-036360-002-original/2024 (H5N1; PQ829557.1), H5HA and N1NA - A/chicken/England/085598/2022 (H5N1; EPI2089022, EPI2089021), human TMPRSS11D/HAT (EAX05559.1) human ST6 beta-galactosamide alpha-2,6-sialyltranferase 1(ST6; AAH31476.1) and CMP-N-acetylneuraminate-beta-galactosamide-alpha-2,3-sialyltransferase 4 (ST3; XP_047283377.1). Mutant versions of these plasmids were generated using site direct mutagenesis. Pseudovirus was produced in HEK 293T cells seeded in 6 well dishes. Cells were co-transfected using lipofectamine 3000 (Invitrogen) at ∼70% confluency with 0.6 µg of the luciferase reporter constructs (pCSFLW), 0.4 µg of the HIV packaging plasmid pGAG-POL, and either 0.4 µg of VSV-G (VSV-G only) or 0.4 µg of the named HA, 0.2 µg of TMPRSS11D (H1 and H7 only) and 0.1 µg of the named NA. Pseudovirus containing supernatants were collected at 48 and 72 hours post-transfection, clarified by centrifugation, aliquoted and frozen at –80°C.

### Pseudovirus receptor binding preference assay

293delST cells were seeded in 6 well plates and transiently transfected at 70% confluency using lipofectamine 3000 with 500 ng of an empty pcDNA3.1 vector, or either ST3 or ST6 expression constructs. 24 hours post-transfection cells were washed, resuspended in fresh media and seeded into 96 well plates at a density of 1 x 10^4 cells per well. Pseudovirus diluted in media containing oseltamivir (to a final concentration of 0.5 µM) was then added to each well. 48 hours post-transduction cells were lysed and read using Bright-Glo reagent (Promega) and a Glomax discover plate reader (Promega). Data for each pseudovirus was normalised to the empty vector control.

### Statistical analysis

GraphPad Prism 10.0 (GraphPad Software, USA) was used for data analysis and visualisation. Ordinary one-way ANOVA Dunnett’s multiple comparisons test and two-way ANOVA multiple comparisons were used to determine statistical significance, where one asterisk indicates 0.01<p<0.05, and 0.001<p<0.01, 0.0001<p<0.001 and p<0.0001 having two, three or four asterisks, respectively. p>0.05 was considered not significant.

### Phylogenetic analysis

All the cattle H5N1 HA sequences (n=4122, last access on 30^th^ June 2025) were acquired from Global Initiative on Sharing All Influenza Data (GISAID) database. The phylogenetic tree was constructed using Molecular Evolutionary Genetics Analysis version 11 (MEGA11) using maximum-likelihood (24). The corresponding amino acid sequences of the H5N1 viruses were analysed using MEGA11. The phylogenetic tree was annotated and visualised using Interactive Tree of Life (iTOL) (25).

### Haemagglutinin structure prediction

H5 HA crystal structure was acquired from Protein Data Bank (https://www.rcsb.org/) with assession number 4JUL. HA structure modelling and prediction were visualised using SWISS-MODEL and visualised and annotated in PyMol version 4.6.

## Results

### Emerging HA mutations in cattle H5N1 viruses

All the available cattle H5N1 virus HA sequences (n=4122) were obtained from GISAID database, and the phylogenetic tree of the shortlisted HA segment was constructed (n=794). Despite most amino acid positions of the HA remaining highly conserved, three amino acid positions: 88, 94, and 131 (H5 mature numbering), are less conserved compared to other HA amino acid positions (Figure 1A). These three amino acids are located in the globular head region of HA, which plays an essential role in virus receptor binding. Of the HA sequences, 65.1% contain D88G, 2.3% contain S94N, and 34.0% contain V131M (Figure 1B). Additionally, we observed polymorphism at position 131, with V131E and V131A. Notably, V131A was also identified in one of the human H5N1 isolates in California (Figure 1C). Among the HA sequences, 94 strains (2.3%) were found to contain the combination of D88G, S94N, and V131M mutations.

**Figure 1.**
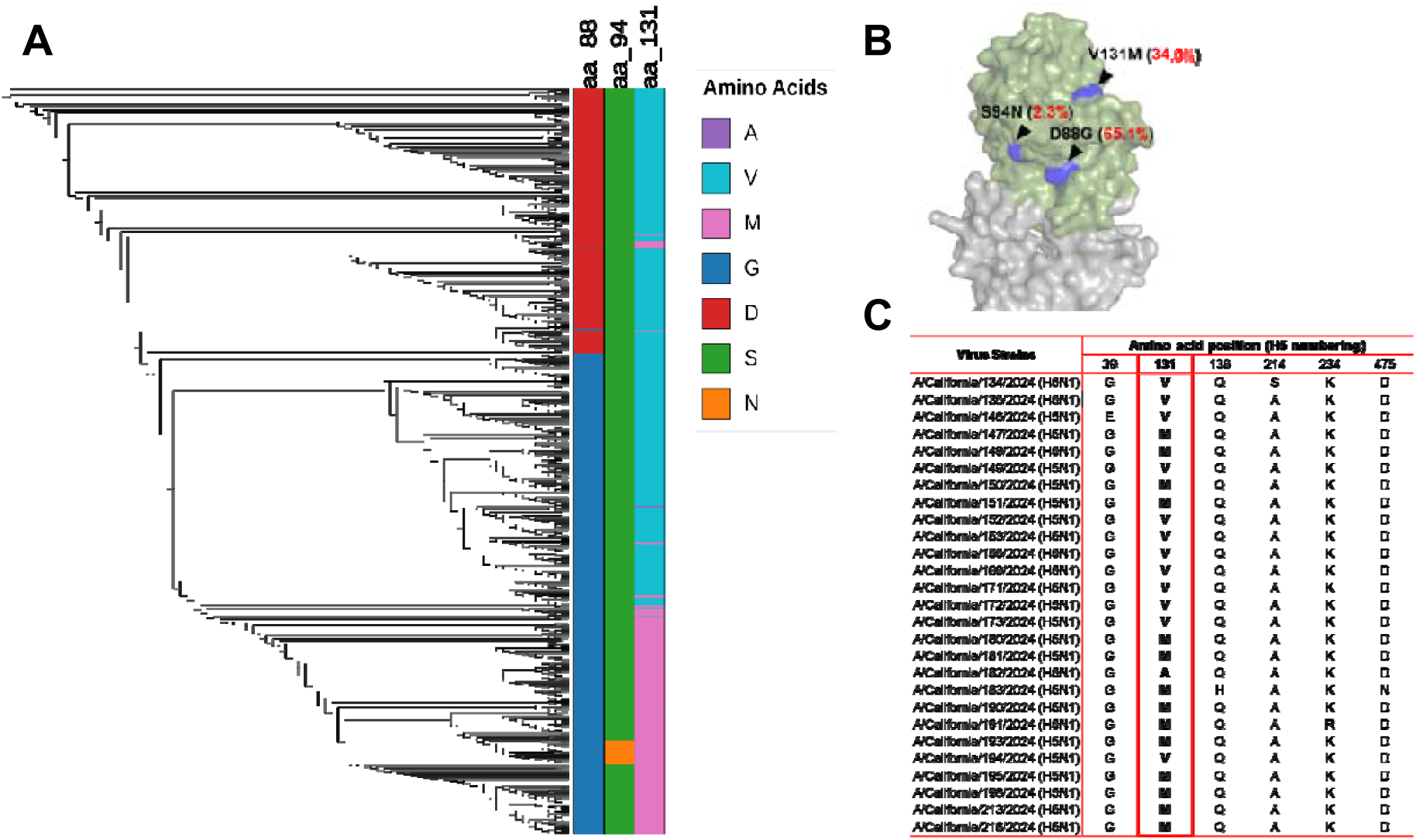
*In silico* analysis of the emerging HA mutations in dairy cattle. (A) Phylogenetic analysis of the cattle H5N1 HA protein sequences (n=794) highlighting the less conserved amino acid positions at 88, 94 and 131 by H5 numbering using available cattle H5N1 sequences on GISAID. (B) Predictive H5N1 HA structure (Assession number 4JUL) highlighting the HA mutations D88G, S94N, and V131M with the frequencies of 65.1%, 2.3%, and 34.0%, respectively. The frequencies are relative to the total cattle H5N1 HA sequences available on GISAID last accessed on 30^th^ June 2025 (n=4122). (C) Amino acid comparison of the HA proteins of human H5N1 in California indicating the polymorphism of 131 position with V131A and V131M.

### The emerging HA mutants from cattle retained binding affinity to avian-like receptors

We first assessed the receptor binding profiles of the precursor cattle wild-type (WT) A/cattle/Texas/24-008749-003/2024 (H5N1) (TX-Cattle WT) and A/cattle/CA/24-036360- 002-original/2024 (H5N1) (CA-Cattle WT), which contains D88G, S94N, and V131M substitutions in combination, using bio-layer interferometry (BLI) with avian-like receptor sugar analogue α-2,3-sialyllactosamine (3SLN) and human-like α2,6-sialyllactosamine (6SLN). Both TX-Cattle WT and CA-Cattle WT showed exclusive binding to 3SLN (Figure 2). A control virus from 2009 H1N1 pandemic A/England/195/2009 (H1N1) showed exclusive binding to human receptor sugar analogue 6SLN (Figure S1).

**Fig. 2.**
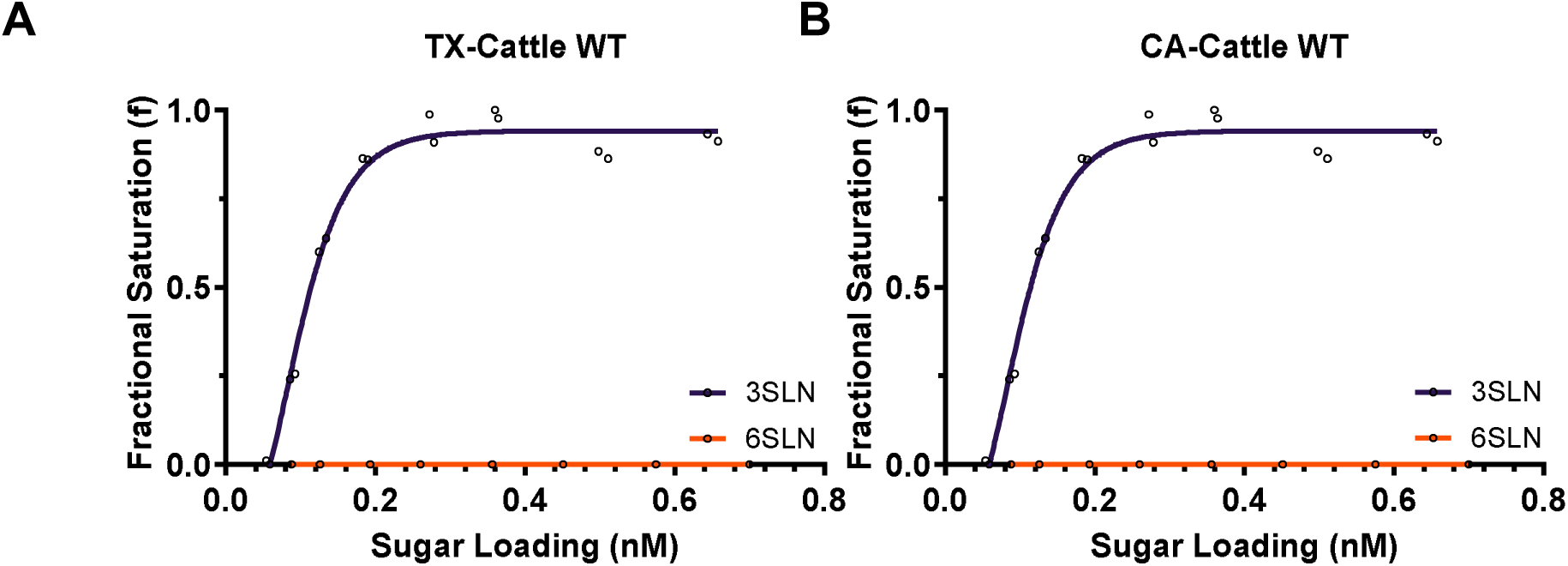
Receptor binding profiles of the precursor H5N1 virus in Texas dairy cattle and the emerging mutant H5N1 virus in California dairy cattle. BLI was used to assess the receptor binding of the H5N1 viruses using avian-like receptor sugar analogue 3SLN and human-like receptor sugar analogue 6SLN. The binding profiles of the (A) Texas dairy cattle H5N1, and (B) California dairy cattle H5N1.

A panel of HA mutant viruses, D88G, S94N, V131A, V131E and V131M, were further generated using TX-Cattle WT HA as a backbone. The panel of mutant viruses also showed binding to avian receptor sugar analogue 3SLN only, with no detectable binding to human receptor sugar analogue 6SLN (Figure 3A-3E). The relative dissociation constant *K_d_* value of 3SLN for each virus was then compared to the TX-Cattle WT, D88G, S94N and V131M showed enhanced binding to 3SLN by 3.5-, 43.6- and 1.2-fold, respectively (Figure 2F). In contrast, V131A and V131E showed decreased binding to 3SLN by 7-fold and 7.5-fold, respectively.

**Figure 3.**
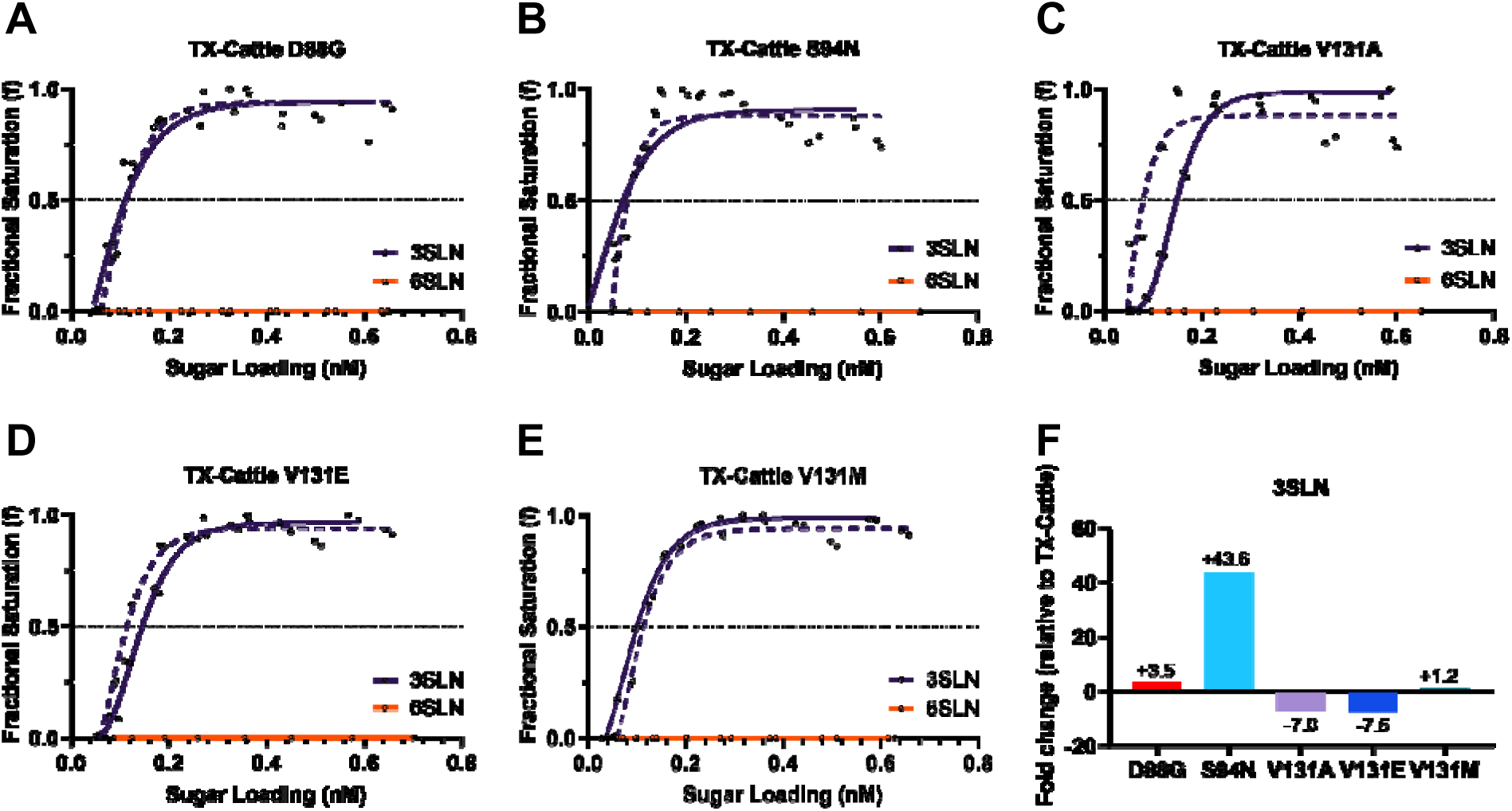
The receptor binding profiles of the emerging HA mutants in dairy cattle. BLI was used to assess the receptor binding of the mutant viruses using avian-like receptor sugar analogue 3SLN and human-like receptor sugar analogue 6SLN. The binding profiles of the mutants (in solid line) were compared to TX-Cattle WT (in dotted line): (A) D88G, (B) S94N, (C) V131A, (D) V131E, (E) V131M. The mutant viruses did not show binding to 6SLN, and the fold change of binding affinity to 3SLN relative to TX-Cattle WT is indicated in (F).

### The emerging HA mutants from cattle retained a preference for avian-type receptors for virus entry

In addition to biophysical interactions between virus and receptor sugar analogues, a pseudovirus approach measuring virus entry was used to further validate the receptor binding results of BLI. HEK293 cells lacking endogenous α2-3 and α2-6 sialyltransferase activities for LacNAc termini on N-glycans, O-glycans and lactoseries glycolipids, were transfected either with an empty vector, an α-2,3-sialyltransferase (ST3GAL4; ST3) or α-2,6- sialyltransferase (ST6GAL1; ST6). This system enabled the assessment of the impact of each mutation in HA mediated cell entry based on luciferase activity as a readout for successful entry quantified by relative light units (RLU). Relative to the empty vector control of each pseudovirus group, TX-Cattle WT, CA-Cattle WT and the TX-Cattle HA mutant viruses (D88G, S94N, V131A, V131E, and V131M) showed enhanced entry exclusively in cells with ST3, but not ST6 overexpressed (Figure 4). Several control pseudoviruses were included in the assay: human pandemic H1N1 showed enhanced entry when ST6, but not when ST3 was overexpressed. Avian H5N1 A/chicken/England/058858/2022 (referred to as AIV48) only showed enhanced entry in cells expressing ST3. However, AIV48 with Q222L mutation (also known as Q226L by H3 numbering), which is known to cause a receptor preference switch from α2,3- to α2,6-linked SA receptors, AIV48 Q222L exhibited enhanced entry into ST6, but not ST3 expressing cells. The HA of the dual-binding H7N9 virus (26), A/Shanghai/02/2013 (H7N9), showed enhanced entry into both cells expressing ST3 and to a lesser extent with ST6.

**Figure 4.**
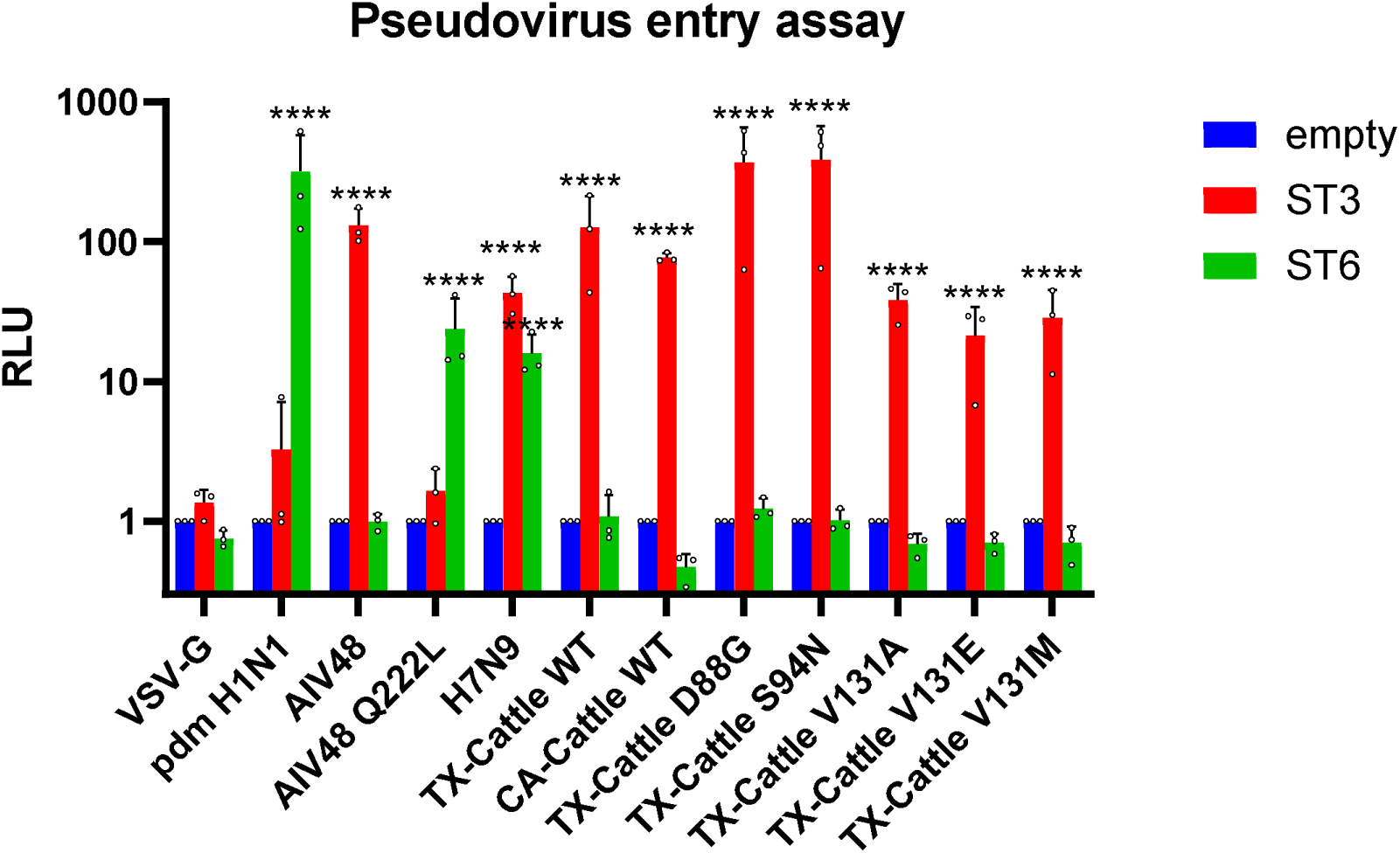
Pseudovirus entry assay for receptor binding preference of the emerging HA mutants in dairy cattle. Monolayered 293delST cells were transiently transfected with pcDNA3.1 empty vector, ST3 or ST6. The cells were then washed, resuspended and seeded into 96-well plates after 24h of transfection. Pseudovirus diluted in culture media supplemented with 0.5 µM oseltamivir was added to each well for 48h. VSV-G served as the negative control, while the pandemic H1N1 A/England/195/2009 (pdmH1N1) virus served as the ST6-positive control. H5N1 AIV48 (A/chicken/England/058858/2022) and its Q222L mutant served as assay controls for ST3- and ST6-mediated entry, respectively. H7N9 (A/Shanghai/02/2013) served as a positive control for entry via both ST3 and ST6. The cells were lysed and read by using Brightglo (Promega), the RLU was measured by microplate reader (Promega). Significance is relative to the empty vector of each pseudovirus. Each point represents a biological repeat from an independent experiment that is the mean of N=3 technical repeats. Data plotted as mean + standard deviation. Statistics performed on log-transformed data (with log-normality confirmed by Shapiro-Wilk test and QQ plot. Significance shown by asterisks indicating: **** P ≤ 0.0001.

### pH stability of the cattle HA mutants

A syncytium formation assay was employed to test the membrane fusion of the WT viruses and panel of single mutant viruses using PBS at different pH values ranging from 5.2 to 6.0. While TX-Cattle WT formed syncytia at pH 5.9, CA-cattle TX fused at a lower pH of 5.6 (Figure 5A). We then assessed the pH fusion of single mutant viruses. TX-Cattle D88G showed membrane fusion at pH 5.6, and HA S94N showed fusion at pH 5.7 (Figure 5B). Both V131A and V131M mutations showed slight increased pH stability with membrane fusion at 5.8. HA mutation with V131E retained the same fusion pH as the WT at pH 5.9. These findings indicate the HA mutations of the cattle H5N1 virus modulate the pH stability, in particular D88G and S94N showed increased pH stability (0.3 and 0.2, respectively) compared to WT virus.

**Fig. 5.**
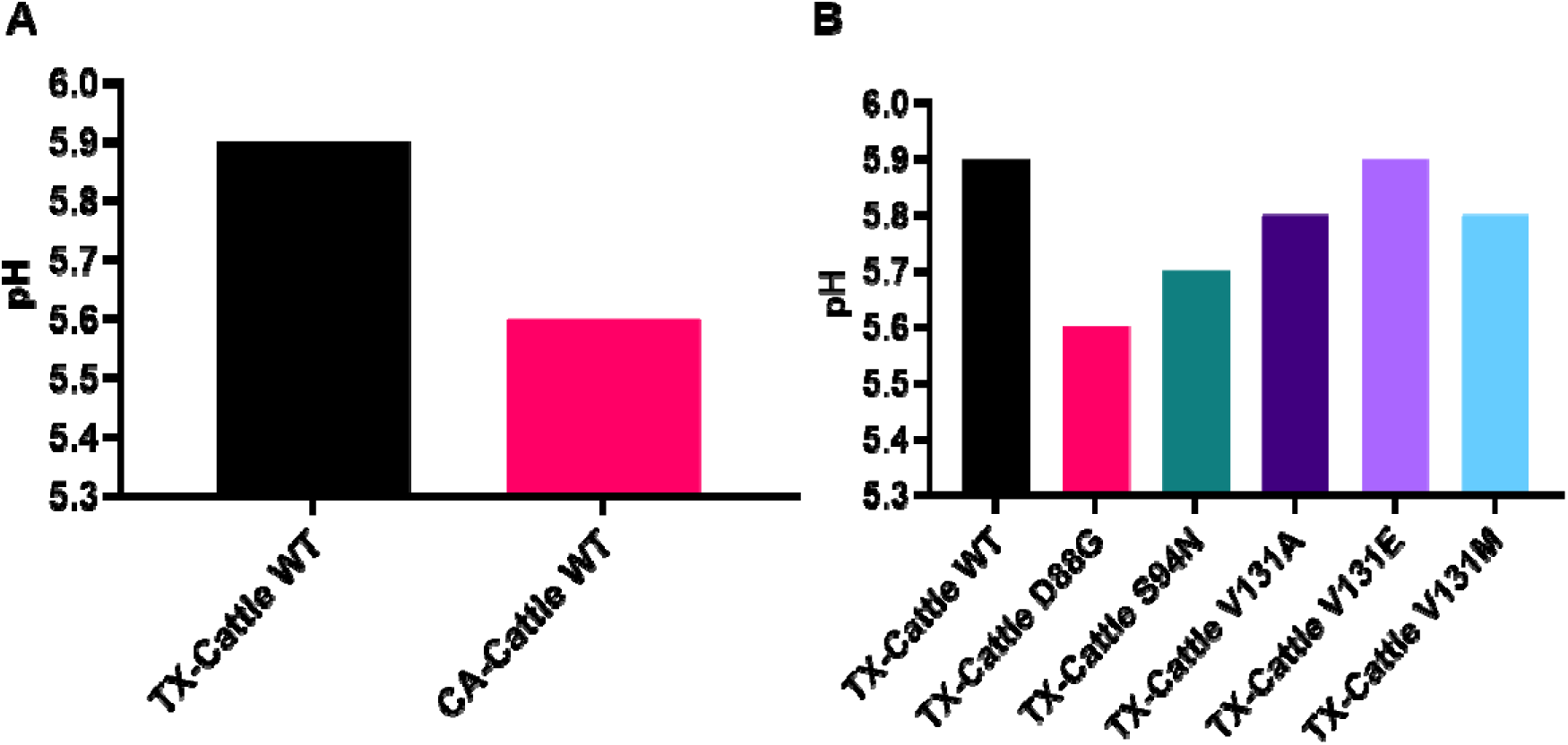
HA stability of the emerging variants in dairy cattle. (A) Comparison of fusion pH between TX-Cattle WT and CA-Cattle WT. (B) Fusion pH of mutant virus with single mutation. Syncytium formation assays in Vero cells were used to assess the fusion pH of the indicated HA mutations and WT. The experiment was repeated twice independently, and the fusion pH was determined as the lowest pH that induced visible syncytium formation.

### Thermostability of the cattle HA mutants

The panel of viruses were heat-treated at different temperatures ranging from 50°C to 60°C for 30 minutes to assess their thermostability, with 4°C used as control. We first compared the thermostability of the precursor TX-Cattle WT and emerging CA-Cattle WT. The thermostability of CA-Cattle WT was significantly increased compared with the TX-Cattle WT virus, with detectable HA activity at 51.9 °C and 53.8 °C (Figure 6A).

**Fig. 6.**
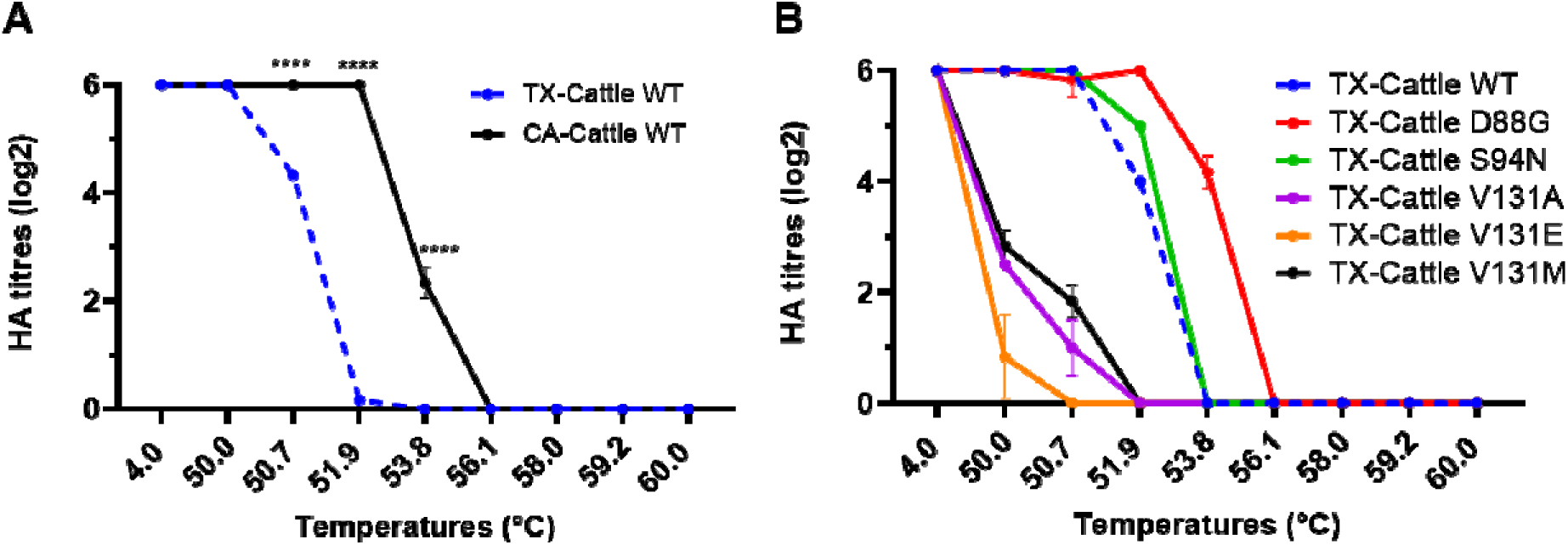
Thermostability of the emerging HA mutants in dairy cattle. (A) Thermostability comparison of precursor TX-Cattle WT and emerging CA-Cattle WT variant. (B) Thermostability of mutant virus with single mutation compared with TX-Cattle WT. The thermostability of the indicated viruses were tested by haemagglutination assay using 1% chicken red blood cells. The titres of the viruses were standardised to 64 HAU and then treated in thermocycler at indicated temperatures for 30min. The experiment was performed three times independently. Significance is relative to TX-Cattle WT. Statistical significance was assessed by two-way ANOVA and indicated by asterisks : ns, not significant (P > 0.05); *, 0.05 ≥ P > 0.01; **, 0.01 ≥ P > 0.001; ***, 0.001 ≥ P > 0.0001; ****, P ≤ 0.0001.

We then assessed viral thermostability of the single mutants. All the tested viruses showed no detectable HA titres above 56.1°C (Figure 5B). Compared to WT, V131A, V131E and V131M showed significant reduction in HAU at 50°C and 50.7°C compared to WT. Statistical significances compared to the WT are listed in Table S2. Furthermore, all three mutations (V131A, V131E, and V131M) showed no detectable HAU at 51.9°C. In contrast, D88G showed increased thermostability at 53.8 °C, and S94N showed marginal increased thermostability at 51.9°C compared to the WT although being statistically significant. Overall, these results suggest that the HA mutations alter the thermostability of the cattle H5N1 virus, with both the D88G and S94N mutations increasing viral thermostability.

## Discussion

Since its emergence in late 2020, the clade 2.3.4.4b HPAIV H5N1 viruses have replaced the precursor H5N8 viruses and rapidly spread worldwide, causing unprecedented losses in the agricultural sector as well as posing significant risk to human health (27). The H5N1 virus has acquired the ability to infect a much broader host range, raising concerns over its heightened zoonotic risk. In particular, mutations in the HA that may increase virus binding affinity to human-like α-2,6 linked SA.

Since the first detection of H5N1 virus in dairy cows in the US, the virus has acquired several mutations in the HA. Several studies have demonstrated cow mammary glands are abundant in both α-2,3 and α-2,6 linked SA (14, 28–30), suggesting that cows may serve as novel mixing vessel for influenza/A viruses and potentially enable the emergence of variants with increased zoonotic potential (31). To assess the receptor binding of the cattle H5N1 viruses, several studies showed that the early bovine-origin H5N1 retained a strong binding preference to avian α-2,3-linked SA (32, 33). Although there was detectable binding to α2,6- linked glycans at a very weak K*_d_* of 56.2µM or >1 mM by surface plasmon resonance (SPR), the binding was considerably weaker than to α2,3-linked glycans (34, 35). We previously assessed the receptor binding profiles of several HA mutations that arose early in the cattle outbreak, in comparison to the early H5N1 strains detected in cattle in Texas. All the tested mutations, including T127A, Q138L, D155N, A156T, P158Q, I162V and Q218R retained binding to avian-like receptors but not to human-like receptors, suggesting the HA of the cattle H5N1 viruses presented a low zoonotic risk (20).

Following the previous study, we further investigated the receptor binding profiles of the emerging HA mutations associated with established infections in cattle and in humans exposed to the cattle-origin H5N1 viruses, in particular we studied viruses in the second epicentre of California where the majority of cases to date have occurred. The tested HA mutations, D88G, S94N, V131A, V131M and V131E, showed continued exclusive binding to an avian-like receptor sugar analogue detected by biolayer interferometry (BLI), bith individually and in combination. To support the BLI data, we also used a functional pseudovirus receptor binding preference assay. Consistent with the BLI results, in HEK293 cells lacking sialyl-transferases, overexpression of ST3, but not ST6 significantly enhanced H5 pseudovirus entry.

The pH stability of HA is one of the key determinants for the route of virus transmission. Oral-fecally spread avian influenza viruses generally have a higher fusion pH (5.6-6.0), in contrast to airborne human influenza viruses that require a lower pH (5.0 to 5.5) (36). Pandemic H1N1 virus with HA H17Y showed more efficient airborne transmissibility in ferrets at pH 5.3 compared to WT at pH 5.5, and the H17Y mutation also showed increased pathogenicity in challenged mice and ferrets (37). The ferret adapted mutation of H3N2 HA P194L reduced fusion pH from 5.7 to 5.2 with increased airborne transmission (38). Similarly, H5N1 HA H103Y changed fusion pH from 5.6 to 5.2, and the mutation also increased virus thermostability as well as airborne transmission in ferrets (17). In this study, the tested mutations, D88G, S94N, V131A, V131E, V131M, and CA-Cattle WT with triple mutations (D88G+S94N+V131M) showed fusion pHs > 5.5, suggesting the mutant viruses retained pH stability consistent with avian viruses.

It is worth noting that over 50% of the bovine H5N1 viruses have now acquired the HA D88G mutation, and D88G showed reduced pH of fusion at 5.6, close to the pH fusion threshold of human viruses at pH 5.5, such as 2009 H1N1 pandemic virus (15, 39). D88G also showed much increased thermostability with detectable HA titre at 53.8°C, suggesting the mutation helps virus to stabilise the HA structure to withstand higher temperatures whilst retaining overall virus fitness for cattle. Interestingly, we noticed that the three mutations at position 131, particularly V131M, reduced viral thermostability compared to TX-Cattle WT, whereas the presence of D88G appeared to partially compensate for the destabilising effect of V131M, as observed in the triple-mutation virus CA-Cattle WT that contains D88G, S94N and V131M. All together highlight a potential role for epistasis in shaping the phenotypic effects of HA mutations in bovine H5N1 viruses.

In conclusion, we assessed the receptor-binding profiles of the emerging bovine-origin H5N1 variants. The mutant viruses tested maintained strong binding to avian-like receptors, with no detectable binding to human-like receptors. The fusion pH of the tested mutations remained above 5.5, suggesting the mutants are unlikely to efficiently transmit between humans. However, mutations D88G and S94N showed higher stability at lower pH and higher temperatures, which is a potential concern given their increasing frequency in cattle origin H5N1 viruses. Considering the widespread nature of cattle farming in the US and the high number of reported infections in dairy cattle, ongoing surveillance and risk assessment of the circulating clade 2.3.4.4b H5N1 viruses, to support risk mitigation measures, remain a high priority for both public health and the agricultural industry.

## Data Availability Statement

All data generated or analysed during this study are included in this manuscript and its supplementary material files.

## Acknowledgement

The authors would like to thank Copenhagen Center for Glycomics at University of Copenhagen for kindly sharing the HEK293 ΔST3GAL1/2/3/4/5/6 ΔST61/2 cells. We thank Dr. Steve Martin for sharing Octet analysis software for receptor binding analysis. We gratefully acknowledge all data contributors, i.e., the authors and their originating laboratories responsible for obtaining the specimens, and their submitting laboratories for generating the genetic sequence and metadata and sharing via the GISAID Initiative, on which this research is based. All submitters of the data may be contacted directly via the GISAID website (https://www.gisaid.org). We gratefully acknowledge U.S. Department of Agriculture (USDA), U.S. Centers for Disease Control and Prevention (CDC), the World Health Organization (WHO) for sharing cattle virus sequences.

---

**Fig. S1.**
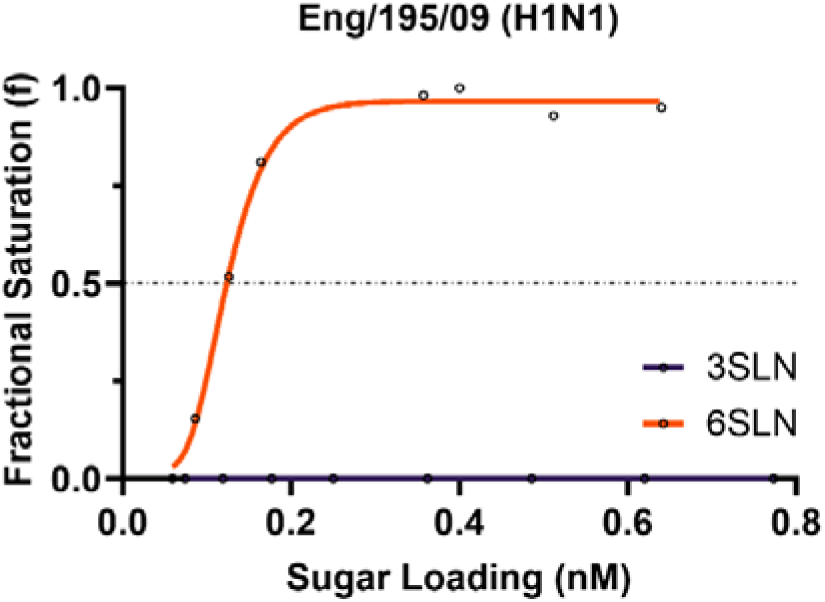
The receptor binding of Eng/195/09 (H1N1). The receptor binding of pandemic H1N1 virus, Eng/195/09, was assessed by BLI using avian-like sugar analogue 3SLN and human-like sugar analogue 6SLN. The virus shows binding affinity to 6SLN and no detectable binding to 3SLN.

**Fig. S2.**
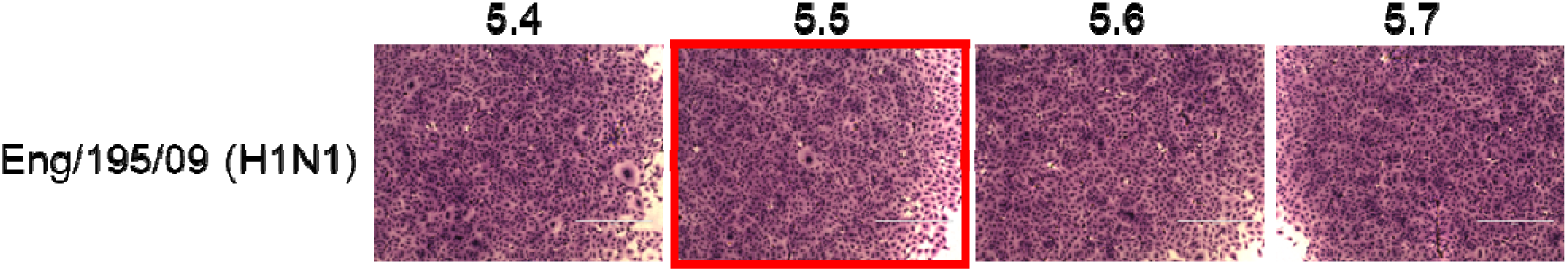
Syncytium formation of Eng/195/09 (H1N1). The lowest pH at which membrane fusion occurs for the pandemic Eng/195/09 (H1N1) is at 5.5.

**Fig. S3.**
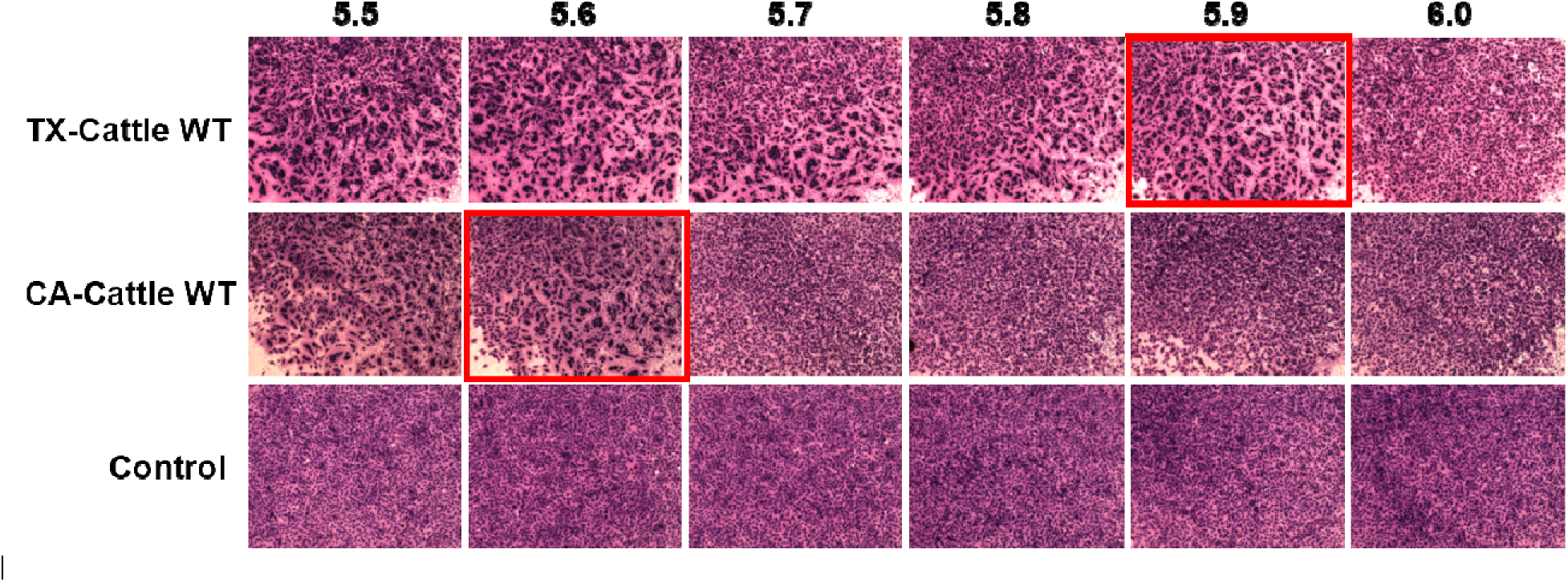
Syncytium formation of the emerging HA mutants in dairy cows. Vero cells were infected with the indicated viruses and treated with PBS at different pH levels. Images highlighted in red indicate the lowest pH at which syncytium formation was observed for each tested virus.

**Fig. S4.**
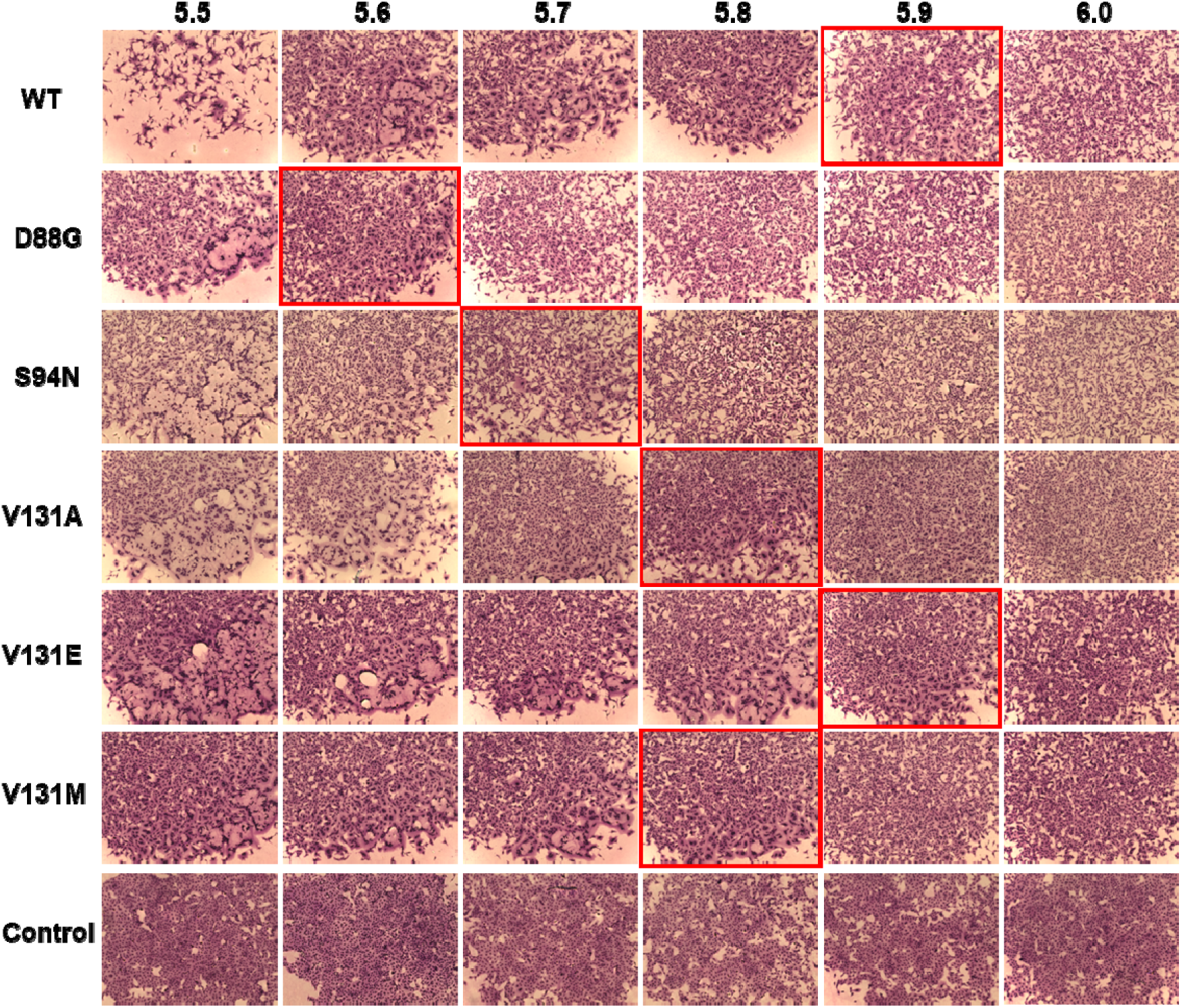
Syncytium formation of the emerging HA mutants in dairy cows. Vero cells were infected with the indicated viruses and treated with PBS at different pH levels. Images highlighted in red indicate the lowest pH at which syncytium formation was observed for each tested virus.

**Table S1.** Estimated relative fold change in Kd values for 3SLN binding by TX-Cattle WT.

| **Virus** | **Estimated $K_d$ values to 3SLN (relative to TX-Cattle WT)** |
| --- | --- |
| **TX-Cattle WT** | **1.00** |
| **D88G** | **3.53** |
| **S94N** | **43.55** |
| **V131A** | **-7.04** |
| **V131E** | **-7.48** |
| **V131M** | **1.22** |

**Table S2.** Statistical analysis of thermostability assay.

| Temperature | Dunnett's multiple comparisons test | Mean Diff. | 95.00% CI of diff. | Below threshold? | Summary | Adjusted P Value |
| --- | --- | --- | --- | --- | --- | --- |
| 4.0 °C | WT vs. D88G | 0 | -0.4015 to 0.4015 | No | ns | >0.9999 |
|  | WT vs. S94N | 0 | -0.4015 to 0.4015 | No | ns | >0.9999 |
|  | WT vs. V131A | 0 | -0.4015 to 0.4015 | No | ns | >0.9999 |
|  | WT vs. V131E | 0 | -0.4015 to 0.4015 | No | ns | >0.9999 |
|  | WT vs. V131M | 0 | -0.4015 to 0.4015 | No | ns | >0.9999 |
| 50.0 °C | WT vs. D88G | 0 | -0.4015 to 0.4015 | No | ns | >0.9999 |
|  | WT vs. S94N | 0 | -0.4015 to 0.4015 | No | ns | >0.9999 |
|  | WT vs. V131A | 3.5 | 3.098 to 3.902 | Yes | **** | <0.0001 |
|  | WT vs. V131E | 5.167 | 4.765 to 5.568 | Yes | **** | <0.0001 |
|  | WT vs. V131M | 3.167 | 2.765 to 3.568 | Yes | **** | <0.0001 |
| 50.7 °C | WT vs. D88G | 0.1667 | -0.2349 to 0.5682 | No | ns | 0.7695 |
|  | WT vs. S94N | 0 | -0.4015 to 0.4015 | No | ns | >0.9999 |
|  | WT vs. V131A | 5 | 4.598 to 5.402 | Yes | **** | <0.0001 |
|  | WT vs. V131E | 6 | 5.598 to 6.402 | Yes | **** | <0.0001 |
|  | WT vs. V131M | 4.167 | 3.765 to 4.568 | Yes | **** | <0.0001 |
| 51.9 °C | WT vs. D88G | -2 | -2.402 to -1.598 | Yes | **** | <0.0001 |
|  | WT vs. S94N | -1 | -1.402 to -0.5985 | Yes | **** | <0.0001 |
|  | WT vs. V131A | 4 | 3.598 to 4.402 | Yes | **** | <0.0001 |
|  | WT vs. V131E | 4 | 3.598 to 4.402 | Yes | **** | <0.0001 |
|  | WT vs. V131M | 4 | 3.598 to 4.402 | Yes | **** | <0.0001 |
| 53.8 °C | WT vs. D88G | -4.167 | -4.568 to -3.765 | Yes | **** | <0.0001 |
|  | WT vs. S94N | 0 | -0.4015 to 0.4015 | No | ns | >0.9999 |
|  | WT vs. V131A | 0 | -0.4015 to 0.4015 | No | ns | >0.9999 |
|  | WT vs. V131E | 0 | -0.4015 to 0.4015 | No | ns | >0.9999 |
|  | WT vs. V131M | 0 | -0.4015 to 0.4015 | No | ns | >0.9999 |
| 56.1 °C | WT vs. D88G | 0 | -0.4015 to 0.4015 | No | ns | >0.9999 |
|  | WT vs. S94N | 0 | -0.4015 to 0.4015 | No | ns | >0.9999 |
|  | WT vs. V131A | 0 | -0.4015 to 0.4015 | No | ns | >0.9999 |
|  | WT vs. V131E | 0 | -0.4015 to 0.4015 | No | ns | >0.9999 |
|  | WT vs. V131M | 0 | -0.4015 to 0.4015 | No | ns | >0.9999 |
| 58.0 °C | WT vs. D88G | 0 | -0.4015 to 0.4015 | No | ns | >0.9999 |
|  | WT vs. S94N | 0 | -0.4015 to 0.4015 | No | ns | >0.9999 |
|  | WT vs. V131A | 0 | -0.4015 to 0.4015 | No | ns | >0.9999 |
|  | WT vs. V131E | 0 | -0.4015 to 0.4015 | No | ns | >0.9999 |
|  | WT vs. V131M | 0 | -0.4015 to 0.4015 | No | ns | >0.9999 |
| 59.2 °C | WT vs. S94N | 0 | -0.4015 to 0.4015 | No | ns | >0.9999 |
|  | WT vs. D88G | 0 | -0.4015 to 0.4015 | No | ns | >0.9999 |
|  | WT vs. V131A | 0 | -0.4015 to 0.4015 | No | ns | >0.9999 |
|  | WT vs. V131E | 0 | -0.4015 to 0.4015 | No | ns | >0.9999 |
|  | WT vs. V131M | 0 | -0.4015 to 0.4015 | No | ns | >0.9999 |
| 60.0 °C | WT vs. D88G | 0 | -0.4015 to 0.4015 | No | ns | >0.9999 |
|  | WT vs. S94N | 0 | -0.4015 to 0.4015 | No | ns | >0.9999 |
|  | WT vs. V131A | 0 | -0.4015 to 0.4015 | No | ns | >0.9999 |
|  | WT vs. V131E | 0 | -0.4015 to 0.4015 | No | ns | >0.9999 |
|  | WT vs. V131M | 0 | -0.4015 to 0.4015 | No | ns | >0.9999 |

## References

1. Lee DH, Bertran K, Kwon JH, Swayne DE. Evolution, global spread, and pathogenicity of highly pathogenic avian influenza H5Nx clade 2.3.4.4. J Vet Sci. 2017;18(S1):269–80.

2. Peacock TP, Moncla L, Dudas G, VanInsberghe D, Sukhova K, Lloyd-Smith JO, et al. The global H5N1 influenza panzootic in mammals. Nature. 2025;637(8045):304–13.

3. Bordes L, Vreman S, Heutink R, Roose M, Venema S, Pritz-Verschuren SBE, et al. Highly Pathogenic Avian Influenza H5N1 Virus Infections in Wild Red Foxes (Vulpes vulpes) Show Neurotropism and Adaptive Virus Mutations. Microbiol Spectr. 2023;11(1):e0286722.

4. Leguia M, Garcia-Glaessner A, Munoz-Saavedra B, Juarez D, Barrera P, Calvo-Mac C, et al. Highly pathogenic avian influenza A (H5N1) in marine mammals and seabirds in Peru. Nat Commun. 2023;14(1):5489.

5. Lindh E, Lounela H, Ikonen N, Kantala T, Savolainen-Kopra C, Kauppinen A, et al. Highly pathogenic avian influenza A(H5N1) virus infection on multiple fur farms in the South and Central Ostrobothnia regions of Finland, July 2023. Euro Surveill. 2023;28(31).

6. Domanska-Blicharz K, Swieton E, Swiatalska A, Monne I, Fusaro A, Tarasiuk K, et al. Outbreak of highly pathogenic avian influenza A(H5N1) clade 2.3.4.4b virus in cats, Poland, June to July 2023. Euro Surveill. 2023;28(31).

7. USDA. Federal and State Veterinary, Public Health Agencies Share Update on HPAI Detection in Kansas, Texas Dairy Herds 2024 [

8. AGRICULTURE USDO. HPAI Confirmed Cases in Livestock | Animal and Plant Health Inspection Service 2025 [Available from: https://www.aphis.usda.gov/livestock-poultry-disease/avian/avian-influenza/hpai-detections/hpai-confirmed-cases-livestock.

9. Pekar JE, Crespo-Bellido A, Lemey P, Bowman AS, Peacock TP, Ochoa JN, et al. Can H5N1 avian influenza in dairy cattle be contained in the US? Cell. 2026;189(3):699–705.

10. CDC. H5N1 Bird Flu Surveillance and Human Monitoring 2025 [Available from: https://www.cdc.gov/bird-flu/h5-monitoring/index.html?cove-tab=1.

11. Organization WH. Avian Influenza A (H5N1)–United States of America. Disease Outbreak News. 2022.

12. Gamblin SJ, Skehel JJ. Influenza hemagglutinin and neuraminidase membrane glycoproteins. J Biol Chem. 2010;285(37):28403–9.

13. Kristensen C, Jensen HE, Trebbien R, Webby RJ, Larsen LE. Avian and Human Influenza A Virus Receptors in Bovine Mammary Gland. Emerg Infect Dis. 2024;30(9):1907–11.

14. Hassard JA, Yang J, Dadonaite B, Pekar JE, Yu J, Richardson SA, et al. Bovine H5N1 influenza viruses have adapted to more efficiently use receptors abundant in cattle. bioRxiv. 2026:2026.04. 02.715584.

15. Peacock TP, Benton DJ, Sadeyen JR, Chang P, Sealy JE, Bryant JE, et al. Variability in H9N2 haemagglutinin receptor-binding preference and the pH of fusion. Emerg Microbes Infect. 2017;6(3):e11.

16. Russell CJ, Hu M, Okda FA. Influenza Hemagglutinin Protein Stability, Activation, and Pandemic Risk. Trends Microbiol. 2018;26(10):841–53.

17. Linster M, van Boheemen S, de Graaf M, Schrauwen EJA, Lexmond P, Manz B, et al. Identification, characterization, and natural selection of mutations driving airborne transmission of A/H5N1 virus. Cell. 2014;157(2):329–39.

18. Narimatsu Y, Joshi HJ, Nason R, Van Coillie J, Karlsson R, Sun L, et al. An Atlas of Human Glycosylation Pathways Enables Display of the Human Glycome by Gene Engineered Cells. Mol Cell. 2019;75(2):394–407 e5.

19. Bull C, Nason R, Sun L, Van Coillie J, Madriz Sorensen D, Moons SJ, et al. Probing the binding specificities of human Siglecs by cell-based glycan arrays. Proc Natl Acad Sci U S A. 2021;118(17).

20. Yang J, Qureshi M, Kolli R, Peacock TP, Sadeyen JR, Carter T, et al. The haemagglutinin gene of bovine-origin H5N1 influenza viruses currently retains receptor-binding and pH-fusion characteristics of avian host phenotype. Emerg Microbes Infect. 2025;14(1):2451052.

21. Xiong X, Xiao H, Martin SR, Coombs PJ, Liu J, Collins PJ, et al. Enhanced human receptor binding by H5 haemagglutinins. Virology. 2014;456-457(100):179–87.

22. Xiong X, Coombs PJ, Martin SR, Liu J, Xiao H, McCauley JW, et al. Receptor binding by a ferret-transmissible H5 avian influenza virus. Nature. 2013;497(7449):392–6.

23. Peacock TP, Goldhill DH, Zhou J, Baillon L, Frise R, Swann OC, et al. The furin cleavage site in the SARS-CoV-2 spike protein is required for transmission in ferrets. Nat Microbiol. 2021;6(7):899–909.

24. Tamura K, Stecher G, Kumar S. MEGA11: Molecular Evolutionary Genetics Analysis Version 11. Mol Biol Evol. 2021;38(7):3022–7.

25. Letunic I, Bork P. Interactive Tree Of Life (iTOL) v5: an online tool for phylogenetic tree display and annotation. Nucleic Acids Res. 2021;49(W1):W293–W6.

26. Belser JA, Gustin KM, Pearce MB, Maines TR, Zeng H, Pappas C, et al. Pathogenesis and transmission of avian influenza A (H7N9) virus in ferrets and mice. Nature. 2013;501(7468):556–9.

27. Cui P, Shi J, Wang C, Zhang Y, Xing X, Kong H, et al. Global dissemination of H5N1 influenza viruses bearing the clade 2.3.4.4b HA gene and biologic analysis of the ones detected in China. Emerg Microbes Infect. 2022;11(1):1693–704.

28. Shi J, Kong H, Cui P, Deng G, Zeng X, Jiang Y, et al. H5N1 virus invades the mammary glands of dairy cattle through ‘mouth-to-teat’ transmission. National Science Review. 2025.

29. Rios Carrasco M, Grone A, van den Brand JMA, de Vries RP. The mammary glands of cows abundantly display receptors for circulating avian H5 viruses. J Virol. 2024;98(11):e0105224.

30. Nelli RK, Harm TA, Siepker C, Groeltz-Thrush JM, Jones B, Twu NC, et al. Sialic Acid Receptor Specificity in Mammary Gland of Dairy Cattle Infected with Highly Pathogenic Avian Influenza A(H5N1) Virus. Emerg Infect Dis. 2024;30(7):1361–73.

31. Pinto RM, Sharp CP, Beeson M, Pankaew N, Hassard JA, Moxom A, et al. The cow udder is a potential mixing vessel for influenza A viruses. bioRxiv. 2025:2025.08. 29.673079.

32. Good MR, Fernandez-Quintero ML, Ji W, Rodriguez AJ, Han J, Ward AB, et al. A single mutation in dairy cow-associated H5N1 viruses increases receptor binding breadth. Nat Commun. 2024;15(1):10768.

33. Santos JJS, Wang S, McBride R, Zhao Y, Paulson JC, Hensley SE. Bovine H5N1 influenza virus binds poorly to human-type sialic acid receptors. bioRxiv. 2024.

34. Song H, Hao T, Han P, Wang H, Zhang X, Li X, et al. Receptor binding, structure, and tissue tropism of cattle-infecting H5N1 avian influenza virus hemagglutinin. Cell. 2025;188(4):919–29 e9.

35. Lin TH, Zhu X, Wang S, Zhang D, McBride R, Yu W, et al. A single mutation in bovine influenza H5N1 hemagglutinin switches specificity to human receptors. Science. 2024;386(6726):1128–34.

36. Russell CJ. Hemagglutinin Stability and Its Impact on Influenza A Virus Infectivity, Pathogenicity, and Transmissibility in Avians, Mice, Swine, Seals, Ferrets, and Humans. Viruses. 2021;13(5).

37. Russier M, Yang G, Rehg JE, Wong SS, Mostafa HH, Fabrizio TP, et al. Molecular requirements for a pandemic influenza virus: An acid-stable hemagglutinin protein. Proc Natl Acad Sci U S A. 2016;113(6):1636–41.

38. Hu M, Kackos C, Banoth B, Ojha CR, Jones JC, Lei S, et al. Hemagglutinin destabilization in H3N2 vaccine reference viruses skews antigenicity and prevents airborne transmission in ferrets. Sci Adv. 2023;9(13):eadf5182.

39. Galloway SE, Reed ML, Russell CJ, Steinhauer DA. Influenza HA subtypes demonstrate divergent phenotypes for cleavage activation and pH of fusion: implications for host range and adaptation. PLoS Pathog. 2013;9(2):e1003151.

